# Dynamic 3D tissue architecture directs BMP morphogen signaling during *Drosophila* wing morphogenesis

**DOI:** 10.1101/411009

**Authors:** Jinghua Gui, Yunxian Huang, Martin Kracklauer, Daniel Toddie-Moore, Kenji Kikushima, Stephanie Nix, Yukitaka Ishimoto, Osamu Shimmi

**Affiliations:** Institute of Biotechnology, University of Helsinki, 00014 Helsinki, Finland; Department of Machine Intelligence and Systems Engineering, Akita Prefectural University, Akita 015–0055, Japan

**Keywords:** Epithelial morphogenesis, three-dimensional architecture, Bone morphogenetic protein (BMP), *Drosophila*, Live-imaging, tissue apposition

## Abstract

At the level of organ formation, tissue morphogenesis drives developmental processes in animals, often involving the rearrangement of two-dimensional (2D) structures into more complex three-dimensional (3D) tissues. These processes can be directed by growth factor signaling pathways. However, little is known about how such morphological changes affect the spatiotemporal distribution of growth factor signaling. Here, using the *Drosophila* pupal wing, we address how Decapentaplegic (Dpp) / Bone Morphogenetic Protein (BMP) signaling and 3D wing morphogenesis are coupled. Dpp, expressed in the longitudinal veins (LVs) of the pupal wing, initially diffuses laterally during the inflation stage to regulate cell proliferation. Dpp localization is then refined to the LVs within each epithelial plane, but with active interplanar signaling for vein patterning, as the two epithelia appose. Our data further suggest that the 3D architecture of the wing epithelia directs the spatial distribution of BMP signaling, revealing how 3D morphogenesis is an emergent property of the interactions between extracellular signaling and tissue shape changes.

## Introduction

Formation of complex 3D tissues from simpler 2D precursors is a basic theme in animal development. 3D architecture formation often involves epithelial morphogenesis, a key process in animal development. Evolutionarily conserved growth factor signaling frequently contributes to these processes. Although how the cellular mechanisms of developmental signaling affect cell and tissue shapes has been actively studied, much less is known about how signaling and dynamic morphogenesis are mutually coordinated (1). Recent advances have indicated how morphogenesis and signaling can be coupled; for example, epithelial structures such as a lumen or villus can regulate the distribution of signaling factors to alter pathway activity (2-4). However, it remains to be addressed how the dynamic 3D tissue architecture affects developmental signaling in a precise spatiotemporal manner.

In vertebrate development, one such example of dynamic 3D architecture formation is tissue fusion, when two apposing tissues approach one another and fuse to form a continuous tissue. This type of process is crucial for the correct formation and functions of many organs and tissues, including the face, neural tube and eyes (5-8). Disruption of fusion leads to various birth defects, including cleft palate, neural tube defects and disorders of eyelid formation (9-11). Although the molecular mechanisms of tissue fusion are likely to be context-dependent, many of the tissue fusion events may share similar mechanisms. Prior to fusion, cellular events such as cell proliferation, apoptosis, migration and epithelial-mesenchymal transition must be coordinated in space and time.

One of the best characterized systems of tissue fusion is the palate, the tissue that separates the oral cavity from the nasal cavity and forms the roof of the mouth. During mammalian embryogenesis, palatogenesis is regulated by a network of signaling molecules and transcription factors to tightly regulate cellular processes (6, 12). Many studies, in both humans and mice, have identified transforming growth factor (TGF)-ß3 as a key signaling factor regulating palatal fusion (13-15). Mice deficient in TGF-ß3 show fully penetrant cleft palate phenotypes, providing an animal model with which to study TGF-ß3 function in palatal fusion (13). TGF-ß3 is expressed in the medial edge epithelial cells prior to adhesion of the opposing palatal shelves, and continues to be expressed during palatal fusion (16). By using a method of palatal shelf organ culture, it has been demonstrated that co-culture of a TGF-ß3 null mutant palatal shelf with wild type palatal shelf resulted in fusion (17). This result suggested that TGF-ß3 produced in wild-type palatal shelf diffused across and rescued the TGF-ß3 mutant shelf, permitting/facilitating fusion.

In *Drosophila*, wing development is a classical model in tissue morphogenesis. The larval wing imaginal disc has been used as a model to address the molecular mechanisms underlying tissue proliferation and patterning. Decapentaplegic (Dpp), a Bone Morphogenetic Protein (BMP) 2/4 type-ligand and member of the TGF-ß family of signaling molecules, has been implicated in regulating a diverse array of developmental events including wing disc development (18). During the larval stage, *dpp* is transcribed in a stripe at the anterior/posterior compartment boundary of the wing imaginal disc, and Dpp forms a long-range morphogen gradient that regulates tissue size and patterning (19, 20). Dpp signaling is needed for tissue proliferation, and Dpp activity gradient formation is crucial for patterning during the late third instar larval stage (21, 22). These processes largely take place in a 2D space, the single cell layer of the wing imaginal disc epithelium.

During the pupal stage that follows, the wing imaginal disc everts to become a two-layered, 3D wing composed of dorsal and ventral epithelial cells (23-26). Previous studies have suggested that pupal wing development is divided into three phases during the first day of pupal development (25, 27, 28). In the first phase, first apposition (0 – 10h after pupariation (AP)), a single-layered wing epithelium everts and forms dorsal and ventral epithelia to become a rudimentary two-layered wing. In the next phase, inflation (10 – 20h AP), the two epithelia physically separate before fusing in the third phase, termed second apposition, at around 20h AP (Figure 1A, Movie S1). During pupal wing development, Dpp signaling is known to play a role in wing vein differentiation. This is largely based on analysis of the *shortvein* group of *dpp* alleles containing deficiencies at the 5’locus that manifest in partial vein loss phenotypes in the adult wing (29, 30). Therefore, dynamic morphological changes in 3D architecture are taking place during the first 24h AP, making this tissue an ideal model to investigate the changes in signaling molecule directionality as a more complex 3D tissue arises from a 2D precursor, and thus how 3D architecture and developmental signaling are coupled.

**Figure 1.**
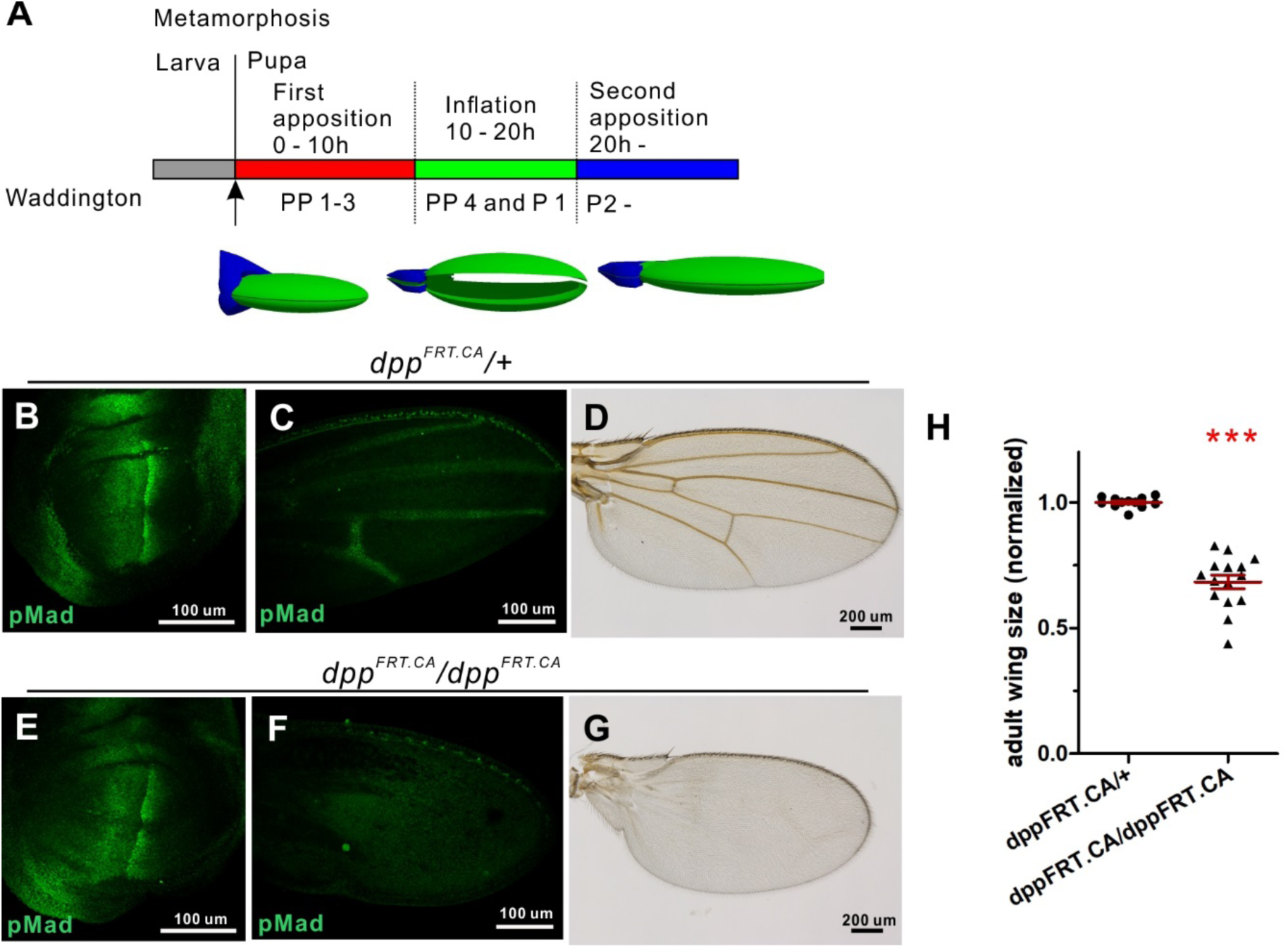
Dpp/BMP signal regulates proliferation and patterning of the *Drosophila* pupal wing. (A) Timing of wing development during the first 24h after pupariation at 25°C. Pupal wing development is divided into three phases; first apposition (0 – 10h AP), inflation (10 – 20h AP), and second apposition (from 20h onwards). Developmental stages (PP (prepupal) 1-4 and P (pupal) 1-2) described by C. H. Waddington are included (27). A schematic of each pupal stage is shown below (hinge in blue and wing in green). Size and tissue shape are not proportional to actual wings. (B-D) pMad staining pattern in wing disc (B), 24h AP pupal wing (C) and an adult wing in control (d*pp*^FRT.CA/+^) (E-G) pMad staining pattern in wing disc (E), 24h AP pupal wing (F), and an adult wing in 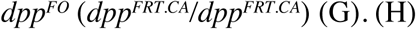 Size comparison between control and *dpp*^*FO*^ wings of adult wings. Means ± SEM, ***P < 0.001, two-paired *t*-test with 95 % confidence intervals (CIs). Larvae were reared at 18°C and then transferred to 29°C 8h before pupariation, followed by dissecting wing imaginal discs (B and E), collecting at 24h AP and dissecting pupal (C and F) or adult stage wings (D and G). Sample sizes: are N = 12 (control) and N = 15 (*dpp*^*FO*^) in (H).

In this study, we re-evaluated the function of Dpp signaling in pupal wing development. Our data reveal that during pupariation, Dpp signaling is needed not only in vein differentiation and patterning, but also has an unexpected key role in tissue proliferation. Specifically, Dpp expressed in the longitudinal veins (LVs) diffuses laterally to regulate tissue size during the inflation stage. N Intriguingly, we find that as dorsal and ventral wing epithelia fuse, the direction of Dpp signaling changes from lateral within each epithelium to interplanar between the epithelia. This results in refinement of Dpp signaling range in the vein regions, which in turn contributes to mirror-image precision of dorsal and ventral epithelial vein patterning. Dpp signaling directionality thus changes from 2D lateral planar to 3D interplanar. Our data further suggest that 3D tissue architecture directs the spatial distribution of Dpp /BMP signaling. These results provide new insights into the mechanism of 3D morphogenesis.

## Results

### Dpp /BMP signal regulates proliferation and patterning of the pupal wing

To re-evaluate the function of Dpp signaling in pupal wing development, we used conditional knockout approaches to remove Dpp in a stage-specific manner. When the knockout was induced in the wing pouch of the wing imaginal disc 24h before pupariation using a conditional *dpp* allele (21), we found that *dpp* expression was efficiently ablated in the pupal wing (Figure S1A, B). Consistent with previous reports, late third instar wing imaginal discs were of equivalent sizes in control and *dpp* knockout animals 24h after induction, even though anti-phosphoMad (pMad) antibody staining, a readout of BMP signaling, was diminished in the wing pouch (Figure S1C, F, I) (21). Intriguingly, pupal wing sizes of *dpp* knockout animals are significantly smaller than in controls at 24h AP, and the BMP signal is lost in *dpp* knockout wings (Figure S1D, G, J). Consistent with this observation, adult wing sizes of *dpp* knockout animals are smaller than that of the control, and wing vein formation is largely abolished (Figure S1E, H, K). Recently, alternative conditional *dpp* knockout alleles have been developed (22), which provide more rapid gene inactivation. Using one of these alternative alleles, we found that BMP signaling was efficiently ablated in the pupal wing, but not in the larval wing imaginal disc, when *dpp* knockout was induced 8h before pupariation (Figure 1B, C, E, F). As shown with the previous knockout allele, these experiments resulted in significantly smaller size and loss of wing vein formation in adult wings compared to controls (Figure 1D, G, H).

To verify independently that these phenotypes are caused by loss of Dpp /BMP signaling in the pupal wing, BMP signal was inhibited in a pupal stage-specific manner by overexpressing *Dad*, an inhibitory Smad (31), resulting both in reduced wing size and in loss of venation in adult wings (Figure S1L, N, P). pMad signaling is also lost in the vein primordia of the pupal wings (Fig. S1M, O). Taken together, these results indicate that the Dpp /BMP signal plays a crucial role in tissue growth and patterning in wing development during pupal stages.

### Growth of the pupal wing involves Dpp/BMP signaling

Positing that Dpp/BMP signaling plays a role in tissue growth of the pupal wing, how is cell proliferation spatiotemporally regulated? Previous studies indicate that cell division in the pupal wing mainly takes place during the inflation stage, without however identifying which molecular mechanisms regulate cell proliferation (32-35). To address whether the Dpp/BMP signal regulates cell proliferation, phosphorylated-histone H3 (pH3) antibody staining was carried out at different time points to detect mitotic cells. The numbers of pH3-positive cells gradually decrease during 18 – 24h AP and are essentially zero at 26h AP in wild-type wings (Figure 2A-F). The numbers of pH3-positive cells in *dpp* knockout wings are significantly lower than control during 18 – 20h AP (Figure 2G-J).

**Figure 2.**
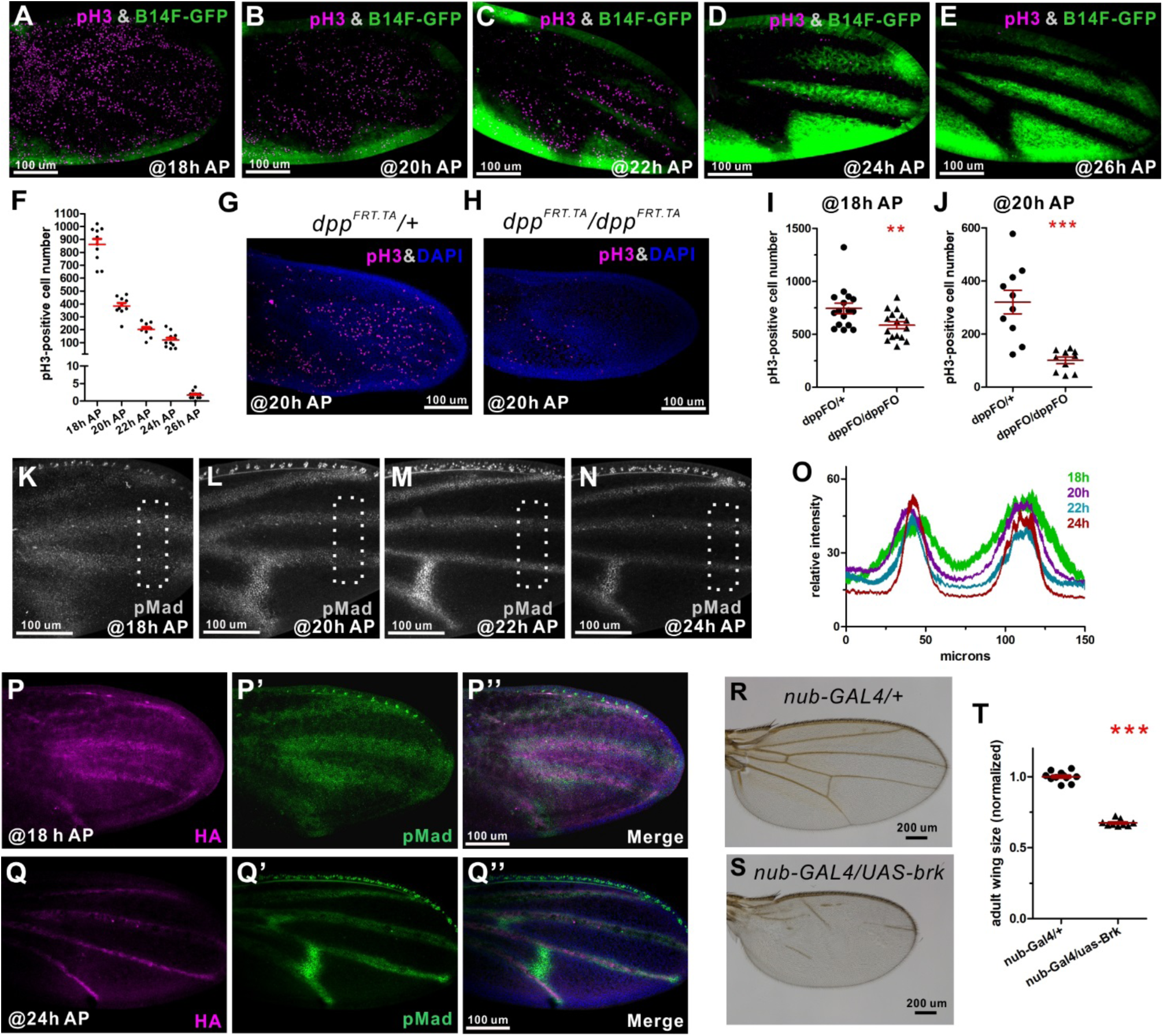
Growth of the *Drosophila* pupal wing involves Dpp/BMP signaling. (A-E) pH3 staining (magenta) and *brk-GFP* (green) of pupal wings in *brk*^*B14F*^*-GFP* in an otherwise wild-type background at 18h (A), 20h (B), 22h (C), 24h (D) and 26h AP (E). (F) Numbers of pH3-positive cells in control at 18h, 20h, 22h, 24h and 26h AP. Mean ± SEM. (G, H) pH3 staining (magenta) and DAPI (blue) in control (G) and conditional knockout (H) at 20h AP. (I, J) Numbers of pH3-positive cells in control (*dpp*^*FRT.TA*^*/+*) and *dpp* conditional knockout (*dpp*^*FRT.TA*^*/dpp*^*FRT.TA*^) at 18h (I) and 20h AP (J). Mean ± SEM. **P < 0.01, ***P < 0.001, two-paired *t*-test with 95 % CIs. (K-N) pMad expression in the pupal wings in wild type at 18h (K), 20h (L), 22h (M), and 24h AP (N). (O) Plot profile analysis of pMad staining in ROIs in (K-N), corresponding to 18h, 20h, 22h and 24h AP. Mean ± SEM, n = 6 for each. (P, Q) Anti-HA antibody staining in *dpp*^*HA*^*/+* animals at 18h (P) and 24h AP pupal wing (Q). pMad expression in the pupal wing (P’, Q’). Merged images of anti-HA (magenta), pMad (green) and DAPI (blue) (P”, Q”). (R, S) Adult wings in control (R) and *brk* overexpression during pupal stage (S). (T) Size comparison between control and *brk* overexpression of adult wings. Larvae were reared at 18°C and then transferred to 29°C after having reached the prepupal stage. Mean ± SEM, ***P < 0.001, two-paired *t*-test with 95 % CIs. Sample sizes are 10 (18h), 10 (20h), 10 (22h), 11 (24h), 10 (26h AP) in (F), 16 (control) and 16 (*dpp*^*FO*^) in (I), 10 (control) and 10 (*dpp*^*FO*^) in (L), and 11 (control) and 9 (*brk* overexpression) in (T).

Since pH3-positive cells are frequently observed in the intervein region, we examined how *dpp* is expressed, and how the Dpp/BMP signal is regulated, during the inflation and second apposition stages. One possibility is that a long-range Dpp signal is needed for cell proliferation during the inflation stage. In order to address this question, we first investigated *dpp* expression in the pupal wing. Similarly to wing imaginal discs, *dpp* is expressed at the anterior-posterior boundary in the early prepupal wing around 5h AP (Figure S2A). Thereafter, expression gradually changes to the positions of future LVs, where it persists until the second apposition stage (Figure S2B-D). We then measured Dpp/BMP signaling by using pMad antibody staining and *brinker (brk)-GFP* (a GFP reporter of the regulatory fragment B14 of *brk*) (36). Brk is a repressor of BMP signal in the wing tissue, expression of which is negatively regulated by BMP signaling (36-39). Our data reveal that the peak level of pMad staining is observed centered on the future LVs, and that lower pMad levels are spread throughout the intervein cells at 18h AP (Figure 2K, O). *brk-GFP* expression is barely detected at the periphery of the pupal wing at 18h AP, indicating that BMP signaling is occurring throughout the pupal wing (Figure 2A) at this time point. When the Dpp/BMP signal was inhibited by overexpressing Dad, pMad expression is not detected, but *brk-GFP* is ubiquitously expressed in the pupal wing at 18h AP (Figure S2E, E’). To address how Dpp ligand is spatiotemporally regulated in the pupal wing, animals expressing HA-tagged Dpp under the control of the genomic *dpp* promoter were utilized (22). HA-Dpp is found not only in the future vein cells (that are ligand-producing cells), but also in intervein cells, at 18h AP (Figure 2P-P”). Taken together, these results suggest that D Dpp forms an activity gradient emanating from future LV cells during the inflation stage.

Intriguingly, the pattern of pH3-positive proliferating cells reflects patterns complementary to *brk* expression (Figure 2A-E). In larval wing imaginal discs, loss of *brk* appears to be sufficient for cell proliferation (40, 41). A recent study further suggests that low-level Dpp signaling (below the level needed for substantial pMad accumulation, but enough for repressing *brk* expression) is sufficient for tissue growth in the wing disc (22). Thus, we examined whether Brk is also a key regulator of proliferation in the pupal wing. Our data reveal that overexpression of *brk* in the wing pouch during the pupal stage resembles loss-of-function of *dpp* in the pupal wing (Figure 2R-T). These results indicate that Dpp trafficking takes place laterally during the inflation stage, and controls cell proliferation by regulating *brk* expression.

As wing development progresses from inflation to second apposition, pMad staining gradually becomes refined to the cells of future LVs, and *brk-GFP* expression is progressively up-regulated in the intervein regions (Figure 2A-E, K-O). Moreover, HA-Dpp is found tightly localized at future vein cells (Figure 2Q-Q”). These results are consistent with previous reports that the Dpp/BMP signal is restricted to LVs at around 24h AP (42). These data further suggest that while the wing tissue is undergoing 3D morphological modifications between the inflation and second apposition stages, the BMP signaling range and pattern are also undergoing dynamic changes.

### Coordination of BMP signaling and patterning between dorsal and ventral epithelia of the pupal wing

What role does Dpp/BMP signaling play in the 3D regulation of growth and patterning in the dorsal and ventral epithelia? First, in order to understand dynamics of 3D structure of the pupal wing, we obtained time-lapse images of optical cross sections of the pupal wing between 18h and 24h AP. These images indicate that maximum distances between dorsal and ventral epithelia are greater than 100 *μ*m at 18h AP, then gradually decrease during re-apposition starting around 20h AP, resulting in dorsal and ventral epithelia of the *Drosophila* wing identical in both size and patterning (i.e. vein and intervein cells) after re-apposition (Movie S2, Figure 3A-F). This raises the question whether patterning and morphogenesis of the two epithelia are regulated independently, or in a coordinated manner. To address this, we next tested how BMP signal transduction acts in wing morphogenesis independently in dorsal and ventral layers. When BMP signaling was reduced only in the dorsal wing epithelium during the pupal stage, by either overexpression of *Dad*, or by knockdown of BMP type-I receptor *thickveins* (*tkv*), adult wings were smaller than in control flies, and displayed partial disruption of vein formation (Figure 3G-I, M). BMP signal transduction was lost in the dorsal pupal wing as expected (Figure 3J-L). Intriguingly, pMad expression in the ventral epithelium was not refined, but instead remained broad at 24h AP, even though the ventral cells are wild-type (Figure 3J’-L’). Furthermore, optical cross sections of the fixed tissues suggest that dorsal and ventral epithelia are not properly fused by 24h AP (Figure 3J”-L”). Taken together, these results indicate that reducing Dpp signaling solely in one 2D epithelial layer nonetheless results in obvious alterations to the 3D structure of the pupal wing, and in reduced size and altered patterning of the final adult wing.

**Figure 3.**
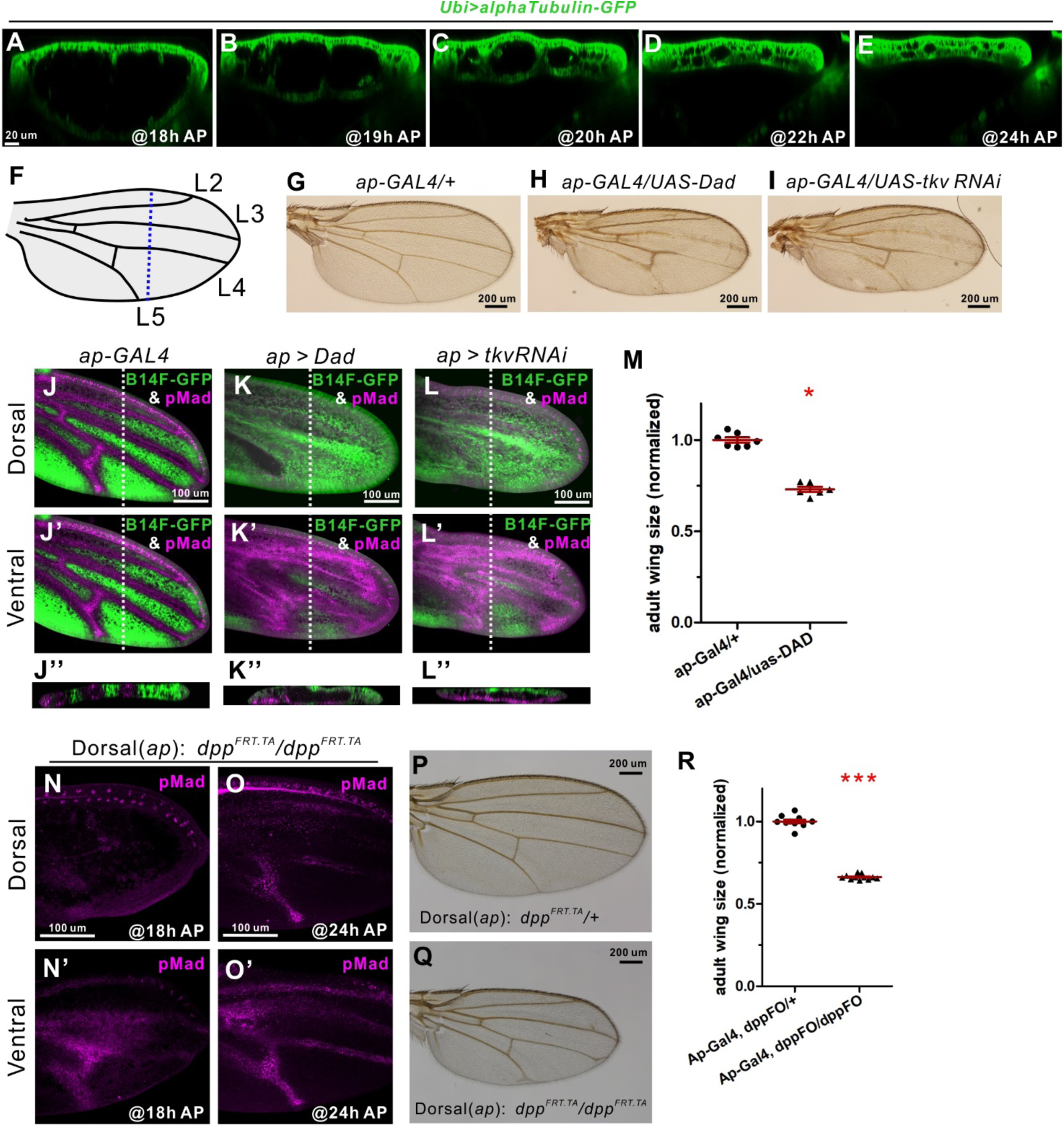
Coordination of BMP signaling between dorsal and ventral epithelia of the pupal wing. (A-E) Time-lapse images of anteroposterior optical cross sections of αTubulin-GFP at 18h (A), 19h (B), 20h (C), 22h (D) and 24h AP (E). Anterior is left and posterior right. Dorsal is up and ventral down. (F) Schematic of pupal wing. Approximate position of imaging in (A-E) is shown as a dotted line. (G-I) Control (G), *Dad* overexpression (H), and *tkv* knockdown adult wings (I). (J-L) pMad expression (magenta) and *brk-GFP* (green) of dorsal (J-L) and ventral tissues (J’-L’) in control (J), *Dad* overexpression (K), and *tkv* knockdown (L) at 24h AP. Optical cross sections focused on the area shown by dotted lines (J”-L”). Dorsal aspect is up, and ventral down. (M) Size comparison between control and *Dad* overexpression adult wings. Larvae were reared at 18° C and then transferred to 29°C after having reached the prepupal stage. Mean ± SEM. *P<0.05, two-paired *t*-test with 95 % CIs. (N, O) pMad expression in dorsal (N, O) and ventral epithelia (N’, O’) in *ap>dpp*^*FO*^ at 18h (N), and at 24h AP (O). (P, Q) adult wings in control (P) and *ap>dpp*^*FO*^ (Q). (R) Size comparison between control and *ap>dpp*^*FO*^ adult wings. Larvae were reared at 18°C and then transferred to 29°C 24h before pupariation, followed by collecting at 18h (N) and 24h AP (O) and dissecting pupal or adult wings (P, Q). Mean ± SEM. ***P < 0.001, two-paired *t*-test with 95 % CIs. Sample sizes are 7 (control) and 6 (*Dad* overexpression) in (M), and 11 (control) and 11 (*ap>dpp*^*FO*^) in (R).

We then studied conditional knockout of *dpp* in the dorsal layer only during pupal stages. In control tissues, pMad expression, the downstream readout of Dpp signaling, shows a similar pattern in dorsal and ventral tissues at both 18h and 24h AP (Figure S3A, B). In contrast, in the conditional *dpp* knockout tissues, pMad expression was only observed in the ventral cells at 18h AP, and thereafter detected in the dorsal cells by 24h AP (Figure 3N, O). Intriguingly, wing vein patterning in conditional knockout adult wings appears largely normal, but tissue size is significantly smaller than in control animals (Figure 3P-R). These results suggest that Dpp ligands expressed in the ventral epithelial layer can induce BMP signaling in the dorsal layer after re-apposition to sustain wing vein development, but tissue proliferation during the inflation stage appears to require ligand production in both dorsal and ventral tissues.

### Interplanar BMP signaling between dorsal and ventral epithelia of the pupal wing

How then is Dpp/BMP signaling regulated between the two epithelial layers? To test whether Dpp mobility is regulated vertically between dorsal and ventral epithelia during the second apposition stage, we employed mosaic analysis with a repressible cell marker (MARCM) (43). When GFP: Dpp -expressing clones are induced in the intervein region of only one epithelial sheet, pMad expression is observed not only in these ligand-producing cells, but also in the opposite epithelium as a near-mirror image at 24h AP (Figure 4A-D). In contrast, pMad expression is only detected in the ligand-producing cells and the flanking regions of the clones, but not in the opposite layer, during wing inflation at 18h AP (Figure S4A, B), in support of the notion that Dpp signal transduction takes place vertically after re-apposition. As a previous report reveals that a positive feedback mechanism through BMP signaling is needed to maintain a short-range Dpp /BMP signal in LVs at 24h AP (42), we examined whether a positive feedback mechanism is also crucial for interplanar BMP signaling. When Dad was overexpressed in Dpp -expressing clones in the intervein region of only one epithelial sheet, pMad expression is observed mostly in the flanking regions of the clones in the same plane. In contrast, pMad is observed at the site of the clones in the opposite epithelial layer in both ligand expressing cells and flanking regions (Figure S4C, D). These results suggest that lateral signaling in the same plane is tightly regulated by active retention through positive feedback mechanisms (42), and in contrast, vertical signaling between the two epithelia appears to be regulated by a distinct mechanism.

**Figure 4.**
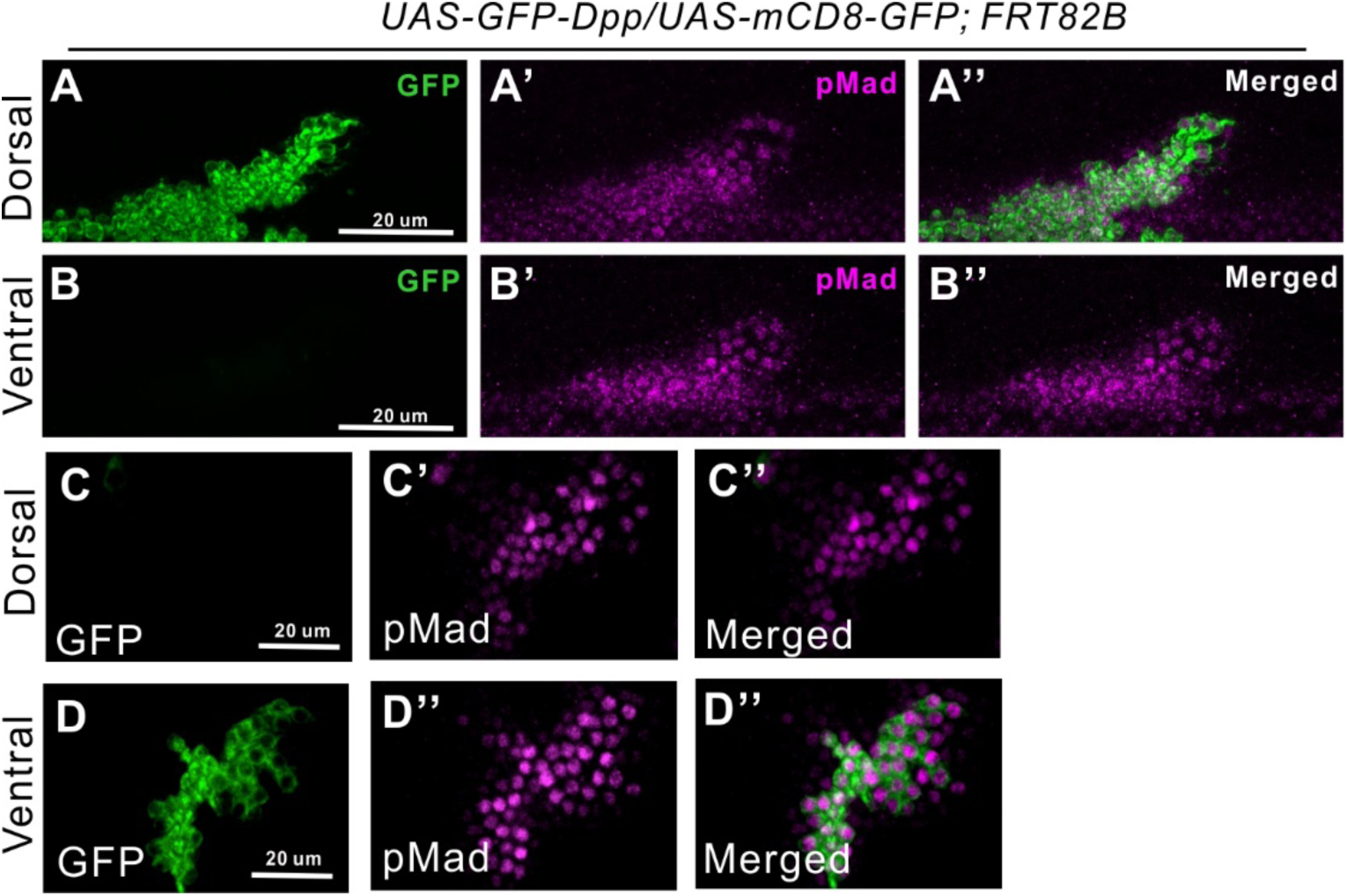
Interplanar BMP signaling between dorsal and ventral epithelia of the pupal wing. (A-D) Dorsal (A, C) and ventral (B, D) epithelia of the pupal wings expressing GFP-Dpp clones in dorsal layer (green, A, B) or ventral layer (green, C, D) at 24h AP. pMad expression (magenta, A’- D’). Merged images (A”-D”).

### Coupling between BMP signaling and 3D tissue architecture

As we observed interplanar signaling between apposed epithelia at 24 h AP, but not during inflation at 18h AP, we asked how interplanar signaling may take place as 3D tissue architecture is formed. One conjecture is that the distance between dorsal and ventral tissues may be a crucial factor in interplanar signaling. The 3D architecture of the developing pupal wing rapidly changes during inflation and second apposition stages (Figure 3A-E, Movie S2). Therefore, we assessed relationships between BMP signaling and 3D architecture of the pupal wing using live time-lapse imaging. Since refinement of BMP signaling can be traced by *brk* expression, we used Brk-GFP flies to obtain time-lapse imaging of optical cross sections of the pupal wing between 18h and 26h AP. RFP-labeled histone H2Av was used to monitor the position of individual cell nuclei (44). Similarly to fixed tissues (Figure 2A-E), Brk-GFP is observed after 20h AP in intervein cells (Movie S3, Figures 5A-F, S5A-F). Importantly, the gap between dorsal and ventral tissues begins to close before *brk* expression is observed. If refinement of BMP signaling in wing vein progenitor cells and 3D tissue architecture are coupled, we expect that 3D tissue dynamics may change when BMP signaling is manipulated. Our data in fixed tissues indicate that 3D architecture is different from control at 24h AP when BMP signaling is disrupted in the dorsal tissues (Figure 3K’’, L’’). To confirm this, we performed live imaging of pupal wings overexpressing Dad in dorsal epithelium. We obtained time-lapse images of optical cross sections of the pupal wing between 18h and 26h AP (Movie S4, Figures 5G-L, S5G-L). Apposition of dorsal and ventral epithelia is significantly delayed, and consequently, Brk-GFP is ubiquitously expressed in the dorsal tissues and less induced in the ventral cells at 24h AP, than in control (Figures 5G-L, S5G-L, Movie S4). These results suggest that the 3D architecture of the pupal wing and spatial distribution of BMP signaling are tightly coupled.

**Figure 5.**
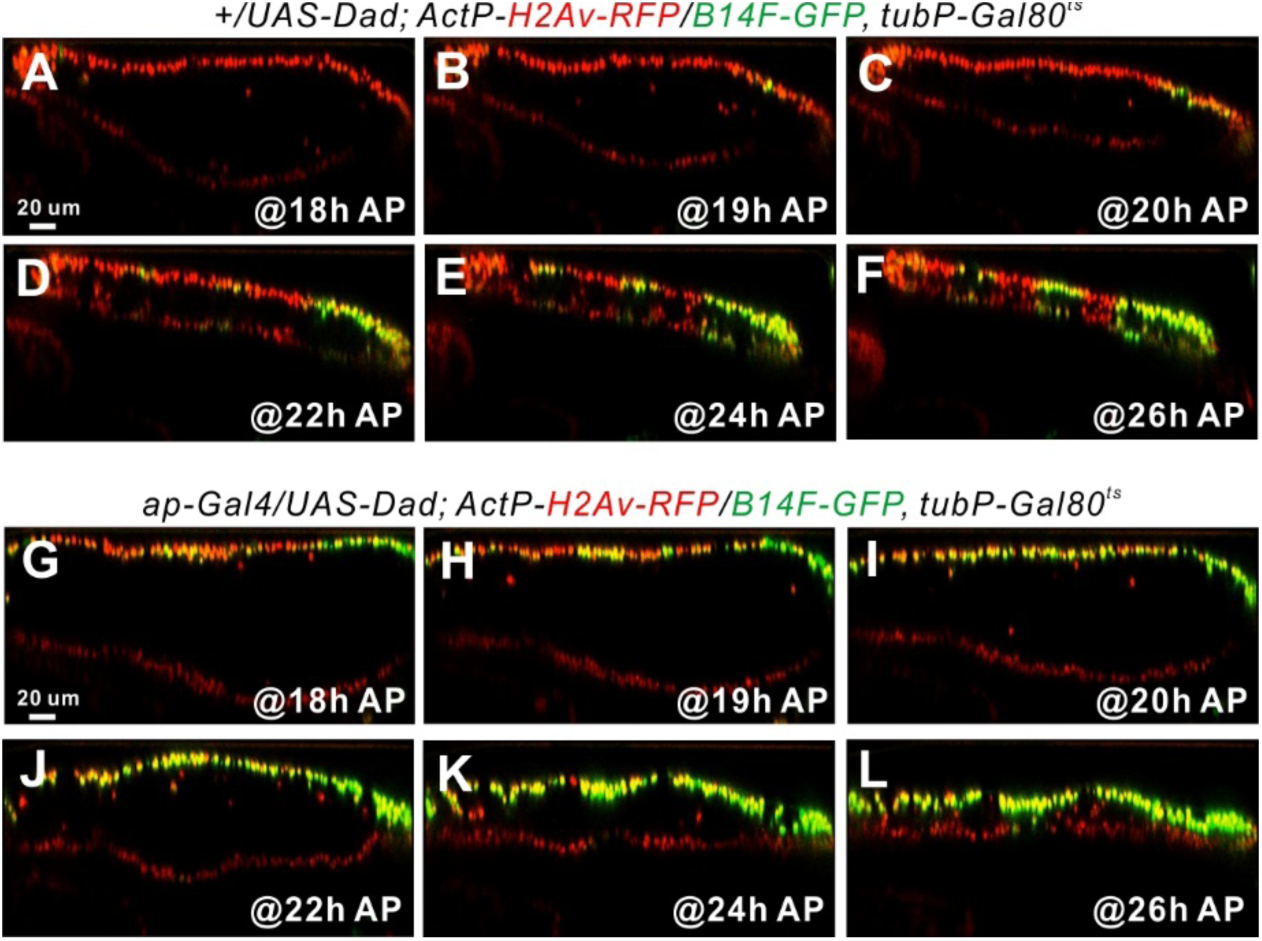
Coupling between BMP signaling and 3D tissue architecture. (A-F) Time-lapse images of anteroposterior optical cross sections in *HistoneH2Av-RFP, brk*^*B14F*^*-GFP* in pupal wing at 18h (A), 19h (B), 20h (C), 22h (D) 24h (E) and 26h AP (F). (G-J) Time-lapse images of anteroposterior optical cross sections of *HistoneH2Av-RFP, brk*^*B14F*^*-GFP* in Dad overexpression in the dorsal epithelium in pupal wing at 18h (G), 19h (H), 20h (I), 22h (J) 24h (K) and 26h AP (L). Anterior is left and posterior right. Dorsal is up and ventral down.

### 3D architecture of the pupal wing instructs spatial distribution of BMP signaling

We next hypothesized that the 3D architecture of wing morphogenesis may be a key regulator of Dpp signaling. To test this idea, we sought to artificially modulate the 3D structure by gently squeezing the pupal abdomen at around 18h AP (Figure 6A, Movies S5, S6). This resulted in excess flow of hemolymph into the wing interepithelial space, causing an increased distance between dorsal and ventral epithelia compared to control animals at 22h AP, and thus extending the inflation stage. Surprisingly, in wings of squeezed pupae, pMad expression is not refined in sharp stripes, and *brk* expression is less induced in the intervein region at 22h AP (Figure 6B, C). Consequently, the proliferation phase appears to last longer, as indicated by more pH3-positive cells at 22h AP than in controls (Figure 6B, D). Importantly, cellular distribution of HA-Dpp is altered with abdominal squeezing. In control tissues, HA-Dpp is highly localized in the future vein cells at basolateral domains (Figure 6E). In contrast, HA-Dpp is dispersed throughout intervein cells in squeezed 24h AP pupal wings (Figure 6F), suggesting that change of 3D architecture affects spatial regulation of Dpp ligands. We also noticed that Tkv distribution, shown by expression of Tkv:YFP, is affected by abdominal squeezing (Figure 6C). Since Tkv levels have been proposed to be a key component in Dpp distribution (45), this further suggests how dynamic changes in 3D tissue structure affect signaling distribution. The effects we observe on HA-Dpp and Tkv:YFP distribution are unlikely to arise due to globally delayed developmental in abdominally squeezed animals, as both squeezed animals and unsqueezed controls develop into adults in a similar time frame (Figure S6). Taken together, these results indicate that 3D architecture of the pupal wing instructs how the Dpp signal is regulated during the processes of tissue proliferation and patterning/differentiation.

**Figure 6.**
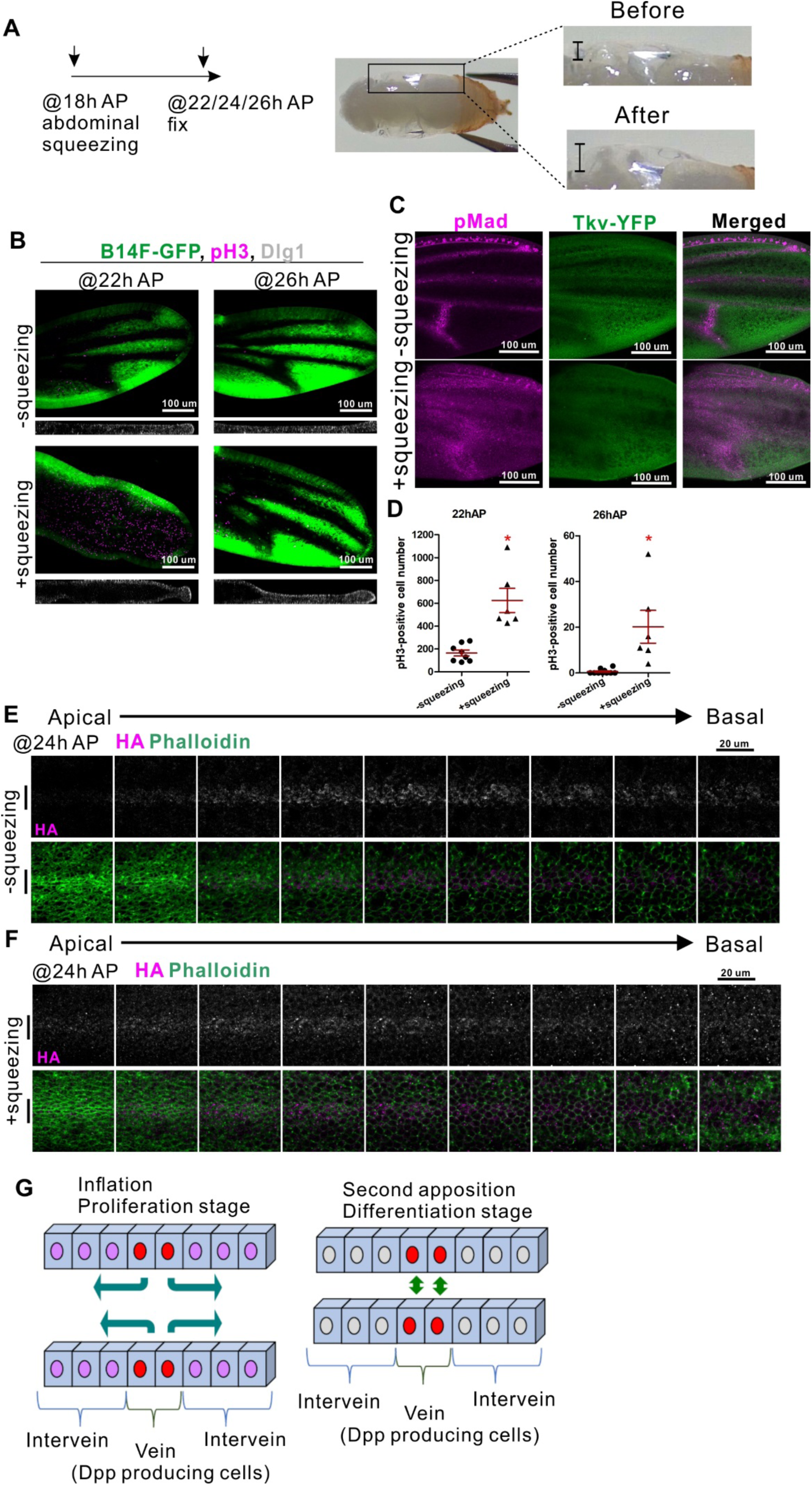
3D architecture of the pupal wing instructs spatial distribution of BMP signaling. (A) Schematic of abdominal squeezing at 18h AP. Pupal tissues were fixed at indicated stages for experiments. Tissue architecture of pupal wings is artificially manipulated through abdominal squeezing (right). Note that inflation between dorsal and ventral epithelia (bars) (wing 3D architecture) is exaggerated after squeezing. (B) *brk-GFP* expression (green) and pH3 staining (magenta) in pupal wings with or without squeezing at 22h and 26h AP. Optical cross sections of the DLG1-stained pupal wings (basolateral staining in wing epithelial cells) are shown in the lower panel. (C) pMad expression (magenta) and Tkv-YFP (green) in pupal wings with or without squeezing at 22h AP. (D) Numbers of pH3-positive cells in pupal wings with or without squeezing at 22h and 26h AP. Mean ± SEM. *P<0.005, two-paired *t*-test with 95 % CIs. Sample sizes are 8 (no squeezing) and 6 (squeezing) at 22h AP, and 10 (no squeezing) and 6 (squeezing) at 26h AP in (D). (E, F) Upper panel: anti-HA antibody staining in pupal wings of *dpp*^*HA*^*/+* without squeezingn(E) or with squeezing (F) at 24h AP. Lower panel: merged images of anti-HA (magenta) and phalloidin (green). Snapshots of 9 different sections of the cells covering L3 along the apicobasal axis at a 1*-μ*m interval. Bars to left of panels show the position of ligand producing cells (future longitudinal vein cells). (G) Schematics of coupling between 3D tissue architecture and Dpp signaling. During the inflation stage Dpp expressed in the longitudinal veins (LVs) diffuses laterally to regulate tissue proliferation. After re-apposition Dpp signaling actively takes place between dorsal and ventral cells to refine signaling range for vein differentiation.

## Discussion

To understand how 3D morphogenesis of an entire tissue and developmental signaling are coordinated, we use the *Drosophila* pupal wing as a model. Historically, although the *Drosophila* pupal wing has not been a widely acknowledged model of 3D tissue architecture formation, the dynamic 3D structure in pupal wing development has been described previously (27), and communication between dorsal and ventral epithelia has also been postulated (46). We propose that the *Drosophila* pupal wing serves as an excellent model for 3D morphogenesis for the following reasons: First, the dynamics of 3D architecture of the pupal wing are observed in a relatively short time period, with three distinct stages, and structural changes and signaling outputs are easily tracked at the cellular level. Second, time-lapse imaging techniques enable us to observe straightforwardly the dynamics of 3D tissue architecture and signaling, and to investigate in real time how morphological changes and signaling are coupled. Third, by using a protocol developed in this study, 3D architecture of the pupal wing can be manipulated without genetic and developmental timing changes (Figures 6, S6, Movies S5, S6). This allows us to investigate experimentally how the assumption of 3D tissue architecture involves spatiotemporal regulation of developmental signaling.

Dpp morphogen signaling in the larval wing imaginal disc has been actively studied as a 2D model (19, 20). Recent studies suggest that Dpp signaling impacts both proliferation and patterning in distinct manners. One study proposes that early stage Dpp signaling is sufficient for tissue proliferation, and the Dpp morphogen gradient at the third instar larval stage is needed for patterning (21). In contrast, a separate study suggests that Dpp signal is needed for proliferation during the third larval instar, at least at a level sufficient to down-regulate *brk* expression (22). In this study, our data suggest that Dpp signaling is required also in the pupal wing for wing cell proliferation and wing vein patterning/differentiation, i.e., well after the third larval instar. This is further highlighted by the fact that these processes are likely affected by the observed changes in Dpp signaling directionality. During the inflation stage, active Dpp trafficking takes place laterally from future LV regions to spread BMP signal throughout the tissue. The pMad staining pattern indicates BMP activity gradient formation centered on LVs (that are ligand producing cells) at 18h AP (Figure 2). It is likely that proliferation rate during pupal wing development is a critical factor to determine final tissue size in the adult. Our data clearly reveal that loss of BMP signal results in reduction of proliferation rate, leading to smaller tissue size at the adult stage. As development progresses from inflation to second apposition, both the pMad staining pattern and *brk* expression reveal that the BMP signaling range becomes refined (Figure 2). Strikingly, BMP signal transduction actively takes place between dorsal and ventral epithelia, playing a crucial role in refinement of the signaling range (Figure 3). These findings suggest that the dynamic interplay between planar and interplanar signaling is linked to coordinate tissue size and patterning.

One of the interesting observations in this study is that tissue size is smaller than control, but overall patterning appears mostly normal, when *dpp* expression was ablated in dorsal cells only (Figure 3P-R). These data clearly support our postulation that Dpp regulates proliferation and patterning/differentiation in distinct manners during pupal wing development. Furthermore, tissue size between dorsal and ventral layers appears to be coordinated when growth signal in only one of the epithelia is manipulated, suggesting the existence of hitherto unidentified mechanisms that coordinate mitosis between dorsal and ventral epithelial cells. Combined with previous studies about Dpp signaling affecting growth and patterning in the larval wing imaginal disc, our data reveal co-optation of the Dpp signaling pathway in the transition from a 2D anlage to a 3D organ.

Our key claim in this work is that formation of 3D tissue architecture and Dpp signaling are tightly coupled (Figure 6G). We support our claim by the following experimental observations. First, changes in the spatiotemporal distribution of Dpp ligand and 3D tissue architecture are coupled. Our data reveal that Dpp ligand distribution changes during inflation and second apposition stages (Figures 2, 4). Importantly, spatial cellular regulation of Dpp ligand appears to be under control of 3D tissue architecture (Figures 5, 6). Second, interplanar signaling between dorsal and ventral cells relies on the distance between the two layered epithelia. Our live-imaging of the pupal wing (Figure 5) and computer simulations (unpublished observations, S.N. and Y.I.) support this claim. This has been further corroborated by changing the 3D architecture of the pupal wing using the abdominal squeezing technique we developed (Figure 6). Importantly, this method simply changes the 3D tissue architecture of the wing without changing genetic background, and does not adversely affect normal developmental timing. Although it remains to be addressed how Dpp ligands move between dorsal and ventral cells, our observations suggest that the basolateral polarity determinant Scribble (Scrib) may be involved in interplanar signaling (Figure S7A, B). Since previous studies showed that Scrib mediates a positive feedback mechanism between BMP signaling and wing vein morphogenesis in the posterior crossvein region of the pupal wing (47), the polarization of epithelial cells may play a role in interplanar signaling. Taken together, we propose that pupal wing morphogenesis and Dpp signaling are coupled, and 3D tissue architecture plays an instructive role in regulating the spatiotemporal distributions of Dpp signaling.

We suspect that mechanisms similar to those found in this study may play roles in the development of many organs and tissues across species. Communication between apposed tissues is likely to be crucial for many developmental processes, but has been insufficiently studied to date. Do cells secrete extrinsic factors to aid the opposing tissues in finding each other across an open space? Is tight coordination of cell proliferation a key process in correct alignment of apposing tissues? If these are the case, what triggers the cellular responses that arise before the tissues come into contact?

In vertebrate embryo development, tissue fusion is observed when two apposing tissues approach one another, and extrinsic factors are often involved in tissue fusion events (5). For example, during formation of the neural tube, which gives rise to the central nervous system, the neuroepithelium forms hinge points and bends on both sides in a U-shape to form the neural folds. The apposed folds approach one another and come into contact to undergo a tissue fusion event that results in formation of the neural tube. Localized BMP signaling plays a crucial role in this process through regulating cellular signaling (48). During eyelid formation, FGF10 produced in the eyelid mesenchyme is required for eyelid fusion and signals in the eyelid epithelium (11). Although it remains to be addressed whether communication between apposed tissues takes place, and how morphological changes and extrinsic factors are coupled, recent advances in mouse embryo *ex vivo* culture techniques have allowed live imaging of these processes (49-51), and hence more dynamic analyses of cellular behaviors during the various fusion processes should be possible.

Furthermore, it is likely that organogenesis from stem cells and tissue self-organization require related mechanisms (52, 53). Characterizing coupling mechanisms between extrinsic signals and morphological changes may therefore further enhance our understanding of organogenesis and morphogenesis.

In summary, our data provide novel insights into how dynamics of 3D tissue architecture instruct spatiotemporal regulation of BMP signaling. We surmise that the concepts highlighted in this work may be generally applicable to molecular mechanisms of animal development, as well as organogenesis from stem cells.

## Acknowledgements

We thank Jukka Jernvall and Irma Thesleff for thoughtful comments on the manuscript. We thank M. Gibson, J. P. Vincent, T. Tabata, C. Gonzalez and G. Pyrowolakis for fly stocks. This work was supported by grant 265648, 308045 from the Academy of Finland, the Sigrid Juselius Foundation, and the Center of Excellence in Experimental and Computational Developmental Biology from the Academy of Finland to OS, JSPS KAKENHI (Grants-in-Aid for Scientific Research) grant number 17K00410 and 17KK0007 to YI, the Integrative Life Science Doctoral Program of the University of Helsinki to JG, and the Finnish Cultural Foundation to YH.

## Author contributions

Conceptualization, J.G. and O.S.; Formal Analysis, S.N. and Y.I.; Investigation, J.G., Y.H., M.K., D.T-M. and O.S.; Writing – Original draft, J.G., M.K. D.T-M. and O.S.; Visualization, K.K.; Supervision, O.S.

1. Gilmour D, Rembold M, & Leptin M (2017) From morphogen to morphogenesis and back. *Nature* 541(7637):311-320.

2. Durdu S, *et al.* (2014) Luminal signalling links cell communication to tissue architecture during organogenesis. *Nature* 515(7525):120-124.

3. Shyer AE, Huycke TR, Lee C, Mahadevan L, & Tabin CJ (2015) Bending gradients: how the intestinal stem cell gets its home. *Cell* 161(3):569-580.

4. Matsuda S, Blanco J, & Shimmi O (2013) A feed-forward loop coupling extracellular BMP transport and morphogenesis in Drosophila wing. *PLoS Genet* 9(3):e1003403.

5. Ray HJ & Niswander L (2012) Mechanisms of tissue fusion during development. *Development* 139(10):1701-1711.

6. Bush JO & Jiang R (2012) Palatogenesis: morphogenetic and molecular mechanisms of secondary palate development. *Development* 139(2):231-243.

7. Nikolopoulou E, Galea GL, Rolo A, Greene ND, & Copp AJ (2017) Neural tube closure: cellular, molecular and biomechanical mechanisms. *Development* 144(4):552-566.

8. Rubinstein TJ, Weber AC, & Traboulsi EI (2016) Molecular biology and genetics of embryonic eyelid development. *Ophthalmic Genet* 37(3):252-259.

9. Dixon MJ, Marazita ML, Beaty TH, & Murray JC (2011) Cleft lip and palate: understanding genetic and environmental influences. *Nat Rev Genet* 12(3):167-178.

10. Greene ND & Copp AJ (2014) Neural tube defects. *Annu Rev Neurosci* 37:221-242.

11. Tao H, *et al.* (2005) A dual role of FGF10 in proliferation and coordinated migration of epithelial leading edge cells during mouse eyelid development. *Development* 132(14):3217- 3230.

12. Smith TM, Lozanoff S, Iyyanar PP, & Nazarali AJ (2012) Molecular signaling along the anterior-posterior axis of early palate development. *Front Physiol* 3:488.

13. Proetzel G, *et al.* (1995) Transforming growth factor-beta 3 is required for secondary palate fusion. *Nat Genet* 11(4):409-414.

14. Kaartinen V, *et al.* (1995) Abnormal lung development and cleft palate in mice lacking TGF-beta 3 indicates defects of epithelial-mesenchymal interaction. *Nat Genet* 11(4):415-421.

15. Iwata J, Parada C, & Chai Y (2011) The mechanism of TGF-beta signaling during palate development. *Oral Dis* 17(8):733-744.

16. Yang LT & Kaartinen V (2007) Tgfb1 expressed in the Tgfb3 locus partially rescues the cleft palate phenotype of Tgfb3 null mutants. *Dev Biol* 312(1):384-395.

17. Taya Y, O‘Kane S, & Ferguson MW (1999) Pathogenesis of cleft palate in TGF-beta3 knockout mice. *Development* 126(17):3869-3879.

18. Shimmi O & Newfeld SJ (2013) New insights into extracellular and post-translational regulation of TGF-beta family signalling pathways. *J Biochem* 154(1):11-19.

19. Affolter M & Basler K (2007) The Decapentaplegic morphogen gradient: from pattern formation to growth regulation. *Nat Rev Genet* 8(9):663-674.

20. Restrepo S, Zartman JJ, & Basler K (2014) Coordination of patterning and growth by the morphogen DPP. *Curr Biol* 24(6):R245-255.

21. Akiyama T & Gibson MC (2015) Decapentaplegic and growth control in the developing Drosophila wing. *Nature* 527(7578):375-378.

22. Sanchez Bosch P, Ziukaite R, Alexandre C, Basler K, & Vincent JB (2017) Dpp controls growth and patterning in Drosophila wing precursors through distinct modes of action. *Elife* 6.

23. Aldaz S, Escudero LM, & Freeman M (2010) Live imaging of Drosophila imaginal disc development. *Proceedings of the National Academy of Sciences of the United States of America* 107(32):14217-14222.

24. Pastor-Pareja JC, Grawe F, Martin-Blanco E, & Garcia-Bellido A (2004) Invasive cell behavior during Drosophila imaginal disc eversion is mediated by the JNK signaling cascade. *Developmental cell* 7(3):387-399.

25. Blair SS (2007) Wing vein patterning in Drosophila and the analysis of intercellular signaling. *Annu Rev Cell Dev Biol* 23:293-319.

26. Matamoro-Vidal A, Salazar-Ciudad I, & Houle D (2015) Making quantitative morphological variation from basic developmental processes: Where are we? The case of the Drosophila wing. *Dev Dyn*.

27. Waddington CH (1940) The genetic control of wing development in Drosophila. *J. Genet.* 41:75-139.

28. Fristrom D, Wilcox M, & Fristrom J (1993) The distribution of PS integrins, laminin A and F-actin during key stages in Drosophila wing development. *Development* 117(2):509-523.

29. Segal D & Gelbart WM (1985) Shortvein, a new component of the decapentaplegic gene complex in Drosophila melanogaster. *Genetics* 109(1):119-143.

30. St Johnston RD, *et al.* (1990) Molecular organization of the decapentaplegic gene in Drosophila melanogaster. *Genes Dev* 4(7):1114-1127.

31. Tsuneizumi K, *et al.* (1997) Daughters against dpp modulates dpp organizing activity in Drosophila wing development. *Nature* 389(6651):627-631.

32. Etournay R, *et al.* (2016) TissueMiner: A multiscale analysis toolkit to quantify how cellular processes create tissue dynamics. *Elife* 5.

33. Milan M, Campuzano S, & Garcia-Bellido A (1996) Cell cycling and patterned cell proliferation in the Drosophila wing during metamorphosis. *Proc Natl Acad Sci U S A* 93(21):11687-11692.

34. O‘Keefe DD, *et al.* (2012) Combinatorial control of temporal gene expression in the Drosophila wing by enhancers and core promoters. *BMC Genomics* 13:498.

35. Schubiger M & Palka J (1987) Changing spatial patterns of DNA replication in the developing wing of Drosophila. *Dev Biol* 123(1):145-153.

36. Muller B, Hartmann B, Pyrowolakis G, Affolter M, & Basler K (2003) Conversion of an extracellular Dpp/BMP morphogen gradient into an inverse transcriptional gradient. *Cell* 113(2):221-233.

37. Campbell G & Tomlinson A (1999) Transducing the Dpp morphogen gradient in the wing of Drosophila: regulation of Dpp targets by brinker. *Cell* 96(4):553-562.

38. Gafner L, *et al.* (2013) Manipulating the sensitivity of signal-induced repression: quantification and consequences of altered brinker gradients. *PLoS One* 8(8):e71224.

39. Minami M, Kinoshita N, Kamoshida Y, Tanimoto H, & Tabata T (1999) brinker is a target of Dpp in Drosophila that negatively regulates Dpp-dependent genes. *Nature* 398(6724):242- 246.

40. Schwank G, Restrepo S, & Basler K (2008) Growth regulation by Dpp: an essential role for Brinker and a non-essential role for graded signaling levels. *Development* 135(24):4003- 4013.

41. Martin FA, Perez-Garijo A, Moreno E, & Morata G (2004) The brinker gradient controls wing growth in Drosophila. *Development* 131(20):4921-4930.

42. Matsuda S & Shimmi O (2012) Directional transport and active retention of Dpp/BMP create wing vein patterns in Drosophila. *Dev Biol* 366(2):153-162.

43. Lee T & Luo L (1999) Mosaic analysis with a repressible cell marker for studies of gene function in neuronal morphogenesis. *Neuron* 22(3):451-461.

44. Schuh M, Lehner CF, & Heidmann S (2007) Incorporation of Drosophila CID/CENP-A and CENP-C into centromeres during early embryonic anaphase. *Curr Biol* 17(3):237-243.

45. Lecuit T & Cohen SM (1998) Dpp receptor levels contribute to shaping the Dpp morphogen gradient in the Drosophila wing imaginal disc. *Development* 125(24):4901-4907.

46. Garcia-Bellido A (1977) Inductive mechanisms in the process of wing vein formation in Drosophila. *Wilhelm Roux’s Arch. Dev. Biol.* 182:93–106.

47. Gui J, Huang Y, & Shimmi O (2016) Scribbled optimizes BMP signaling through its receptor internalization to the Rab5 endosome and promote robust epithelial morphogenesis. *PLoS Genet*.

48. Eom DS, Amarnath S, Fogel JL, & Agarwala S (2011) Bone morphogenetic proteins regulate neural tube closure by interacting with the apicobasal polarity pathway. *Development* 138(15):3179-3188.

49. Massarwa R & Niswander L (2013) In toto live imaging of mouse morphogenesis and new insights into neural tube closure. *Development* 140(1):226-236.

50. Williams M, Yen W, Lu X, & Sutherland A (2014) Distinct apical and basolateral mechanisms drive planar cell polarity-dependent convergent extension of the mouse neural plate. *Dev Cell* 29(1):34-46.

51. Yamaguchi Y, *et al.* (2011) Live imaging of apoptosis in a novel transgenic mouse highlights its role in neural tube closure. *J Cell Biol* 195(6):1047-1060.

52. Lancaster MA & Knoblich JA (2014) Organogenesis in a dish: modeling development and disease using organoid technologies. *Science* 345(6194):1247125.

53. Sasai Y (2013) Cytosystems dynamics in self-organization of tissue architecture. *Nature* 493(7432):318-326.

54. Buttitta LA, Katzaroff AJ, Perez CL, de la Cruz A, & Edgar BA (2007) A double-assurance mechanism controls cell cycle exit upon terminal differentiation in Drosophila. *Developmental cell* 12(4):631-643.

55. Classen AK, Aigouy B, Giangrande A, & Eaton S (2008) Imaging Drosophila pupal wing morphogenesis. *Methods Mol Biol* 420:265-275.

## Materials and Methods

### Fly genetics

*nub-GAL4* (#25754), *ap-GAL4* (#3041), *tubP-GAL80*^*ts*^ (#7017), w; *UAS-GFP-dpp* (#53716) *w; His2Av-RFP* (#23650), *w;; His2Av-RFP* (#23651), and *scrib*^2^ (#41775) were obtained from the Bloomington *Drosophila* Stock Center. *UAS-tkv* RNAi (#3059) was obtained from the Vienna *Drosophila* RNAi Center. *Tkv-YFP*^*CPTI00248*^ (#115298) was obtained from the Kyoto *Drosophila* Genetic Resources Center. *dpp*^*FRT.CA*^*; rn-Gal4 and dpp*^*FRT.CA*^, *UAS-Flp were obtained from* JP Vincent, *dpp*^*FO*^*; UAS-FLP, dpp*^*FO*^ *nub-GAL4, and dpp*^*FO*^ *ap-Gal4 (dpp*^*FO*^ named as *dpp*^*TA*^ in this paper) from M. Gibson, *w, alphaTubulin-GFP from* C. Gonzalez, *UAS-Dad* from T. Tabata, and *brk*^*B14F*^*-GFP and UAS-brk* from G. Pyrowolakis. Fly stocks were maintained at 25°C unless otherwise mentioned.

To generate the *dpp*^*FO*^ mutant, mid-3rd instar larvae, raised at 1°C for 7-8 days after egg laying (AEL), were shifted to 2° C for 8h (*dpp*^*FRT.CA*^) (22) or 24h before pupariation (*dpp*^*FRT.TA*^) (21). Then the late-3rd instar larvae and white prepupae were subjected to the subsequent experiments, including immunostaining and in situ hybridization.

For exogenous expression of *Dad* or shRNA at pupal stages, white prepupae raised at 18° C or room temperature were shifted to 29° C, and the pupae of indicated ages were collected and subjected to the subsequent experiments.

For MARCM analysis, flies were maintained at 25° C throughout development, except for heat-shock treatment. Three days AEL, 2nd instar larvae underwent heat shock for two hours in a 37°C water bath. Thereafter, white prepupae were collected, and those aged to 24 hours were fixed and subjected to immunostaining analysis.

Pupal wings were dissected at developmental timepoints equivalent to 25°C. Calculations for developmental timing at 2° C were based on previously published data (54).

### Full genotypes

Figure 1B-D: *w; dpp*^*FRT.CA*^, *UAS-Flp/+; rn-Gal4/tubP-Gal80*^*ts*^

Figure 1E-G: *w; dpp*^*FRT.CA*^, *UAS-Flp/dpp*^*FRT.CA*^*; rn-Gal4/tubP-Gal80*^*ts*^

Figures 2A-E, 6b: *brk*^*B14F*^*-GFP* (III)

Figures 2G: *w; nub-Gal4/dpp*^*FRT.TA;*^ *UAS-Flp/tubP-Gal80*^*ts*^

Figures 2H: *w; dpp*^*FRT.TA*^, *nub-Gal4/dpp*^*FRT.TA;*^

*UAS-Flp/tubP-Gal80*^*ts*^

Figures 2K-N, 6C: *w; Tkv-YFP*^*CPTI00248/*^*+*

Figures 2P, Q, 6E, F: *w; dpp*^*FRT.CA/*^*+*

Figure 2R: *w; nub-Gal4*/*UAS-brk*; *tubP-Gal80*^*ts*^*/+*

Figure 2S: *w; nub-Gal4*/*+*; *tubP-Gal80ts/+*

Figure 3A-E: *w, ubi-aTubulin:GFP*

Figure 3G: *ap-Gal4*/*+*; *tubP-Gal80*^*ts*^*/+*

Figure 3H: *w; ap-Gal4*/*UAS-Dad*; *tubP-Gal80*^*ts*^*/+*

Figure 3I: *w; ap-Gal4*/*UAS-tkv*^*RNAi*^; *tubP-Gal80*^*ts*^*/+*

Figure 3J: *ap-Gal4*/*+*; *tubP-Gal80ts brk*^*B14F*^*-GFP/+*

Figure 3K: *w; ap-Gal4*/*UAS-Dad*; *tubP-Gal80*^*ts*^ *brk*^*B14F*^*-GFP/+*

Figure 3L: *w; ap-Gal4*/*UAS-tkv*^*RNAi*^; *tubP-Gal80*^*ts*^ *brkB14F-GFP/+*

Figures 3N, O, Q: *w; dpp*^*FRT.TA*^, *ap-Gal4/d*^*ppFRT.TA*^*; UAS-Flp/tubP-Gal80*^*ts*^

Figures 3P: *w; ap-Gal4/dpp*^*FRT.TA*^*; UAS-Flp/tubP-Gal80*^*ts*^

Figure 4A-D: *hs-Flp; tubP-Gal4 UAS-mCD8-GFP/UAS-GFP-dpp; tubP-Gal80*^*ts*^ *FRT82B/FRT82B*

Figure 5A-F: *w; +/UAS-Dad; His2Av-RFP/brk*^*B14F*^*-GFP, tubP-Gal80*^*ts*^

Figure 5G-L: *w; ap-Gal4/UAS-Dad; His2Av-RFP/brk*^*B14F*^*-GFP, tubP-Gal80*^*ts*^

Figure S1A, F-H: *w; dpp*^*FRT.TA,*^ *nub-Gal4/dpp*^*FRT.TA*^*; UAS-Flp/tubP-Gal80*^*ts*^

Figure S1B-E: *w; nub-Gal4/dpp*^*FRT.TA*^*; UAS-Flp/tubP-Gal80*^*ts*^

Figure S1L, M: *w; nub-Gal4*/*+*; *tubP-Gal80*^*ts*^*/+*

Figure S1N, O: *w; nub-Gal4*/*UAS-Dad*; *tubP-Gal80*^*ts*^*/+*

Figure S2A-D: *yw*

Figure S2E: *w; nub-Gal4*/*UAS-Dad*; *tubP-Gal80*^*ts*^*/brk*^*B14F*^*-GFP*

Figures S3A, B: *w; ap-Gal4/dp*^*pFRT.TA*^*; UAS-Flp/tubP-Gal80*^*ts*^

Figure S4A, B: *hs-Flp; tubP-Gal4 UAS-mCD8-GFP/UAS-GFP-dpp; tubP-Gal80*^*ts*^*FRT*^82B^*/FRT*^*82B*^

Figure 4C, D: *hs-Flp tubP-Gal80 FRT*^*19A*^*/FRT*^*19A*^*; tubP-Gal4 UAS-mCD8-GFP/UAS-Dad; UAS-GFP-dpp/+*

Figure S5A-F: *w; +/UAS-Dad; His2Av-RFP/brk*^*B14F*^*-GFP, tubP-Gal80*^*ts*^

Figure S5G-L: *w; ap-Gal4/UAS-Dad; His2Av-RFP/brkB14F-GFP, tubP-Gal80ts*

Figure S6: *Oregon R*

Figure S7A, B: *hs-Flp; tubP-Gal4 UAS-mCD8-GFP/UAS-GFP-dpp; tubP-Gal80*^*ts*^ *FRT*^*82B*^*/FRT*^*82B*^*scrib*^2^

### Immunohistochemistry

Pupae were fixed in 3.7% formaldehyde (Sigma-Aldrich®) at 4°C overnight, after which pupal wings were dissected. Larvae were fixed in 3.7% formaldehyde at room temperature for 20 minutes, after which wing imaginal discs were dissected. The following primary antibodies were used: mouse anti-DLG1 (1:50; Developmental Studies Hybridoma Bank (DSHB), University of Iowa), rabbit anti-phospho-SMAD1/5 (1:300; Cell Signaling Technologies), rabbit anti-phospho-Histone H3 (1:500; Millipore), rat anti-HA 3F10 (1:100; Roche). Alexa 488 conjugated phalloidin (1:200; Thermo Fisher Scientific). Secondary antibodies were anti-mouse IgG Alexa 488, anti-rabbit IgG Alexa 568, anti-mouse IgG Alexa 647, and-rat IgG Alexa 568 (1:200; Thermo Fisher Scientific).

### Imaging and image analysis

Fluorescent images were obtained with a Zeiss LSM700 upright laser confocal microscope, in situ hybridization images and adult wing images were obtained with a Nikon ECLIPSE 90i microscope. All images were processed and analyzed with ImageJ (NIH) software. Images subjected to intensity measurement were captured with the same parameters, otherwise, the images were adjusted with linear methods. The “remove outliner” function of ImageJ was applied to remove bright speckles, which are non-specific signal, in some images. None of the processing steps affect data interpretation.

### Time lapse imaging

Prepupae of indicated genotypes were raised and collected at room temperature, then shifted to 29°C until staged appropriately (late inflation stage, roughly 18h AP equivalent at 25 °C). Pupae were retrieved, briefly rinsed in water, dried on a Kimwipe, then positioned on a piece of double-sided tape (right wing facing up). Windows were carefully dissected into the pupal cases in the region of the wing using a micro-knife (Fine Science Tools, cat# 10316-14) essentially as described (55), avoiding damage to the underlying tissue. A tiny drop of halocarbon oil (Sigma Aldrich) was applied to the exposed pupal wing with a disposable pipet tip to prevent tissue desiccation during imaging. The pupae, adhering to strips of double-sided tape cut with a disposable scalpel, were then placed oil-side down onto a 24 × 50 mm coverslip. After 5-6 pupae were collected onto the coverslip, wings were time-lapse imaged on a Leica SP8 STED confocal microscope by taking optical anteroposterior cross sections of each wing every 4-5 minutes using the xzyt-function. The resulting time lapse images were processed into AVI-format videos using Imaris v.9.1.2 (Bitplane/Oxford Instruments).

### Modulation of tissue architecture of the pupal wings through abdominal squeezing

Pupal cases of 18h AP pupae were carefully removed from the anterior to expose the wings. Then, the pupae were positioned with dorsal side facing up before forceps were used to clasp the abdomen. Once the abdomen was stabilized, we exerted force by gently squeezing the abdomen with the forceps. The force was gradually increased until influx of hemolymph into the wings was observed, which causes the enhanced inflation. After squeezing, the pupae were maintained at 25°C in a humid chamber and subjected to the experiments at the indicated time points.

### Statistics

All experiments were carried out independently at least three times. Data are means ± 95 % confidence intervals (CIs). Statistical significance was calculated by the two paired *t*-test method.

---

**Figure S1.**
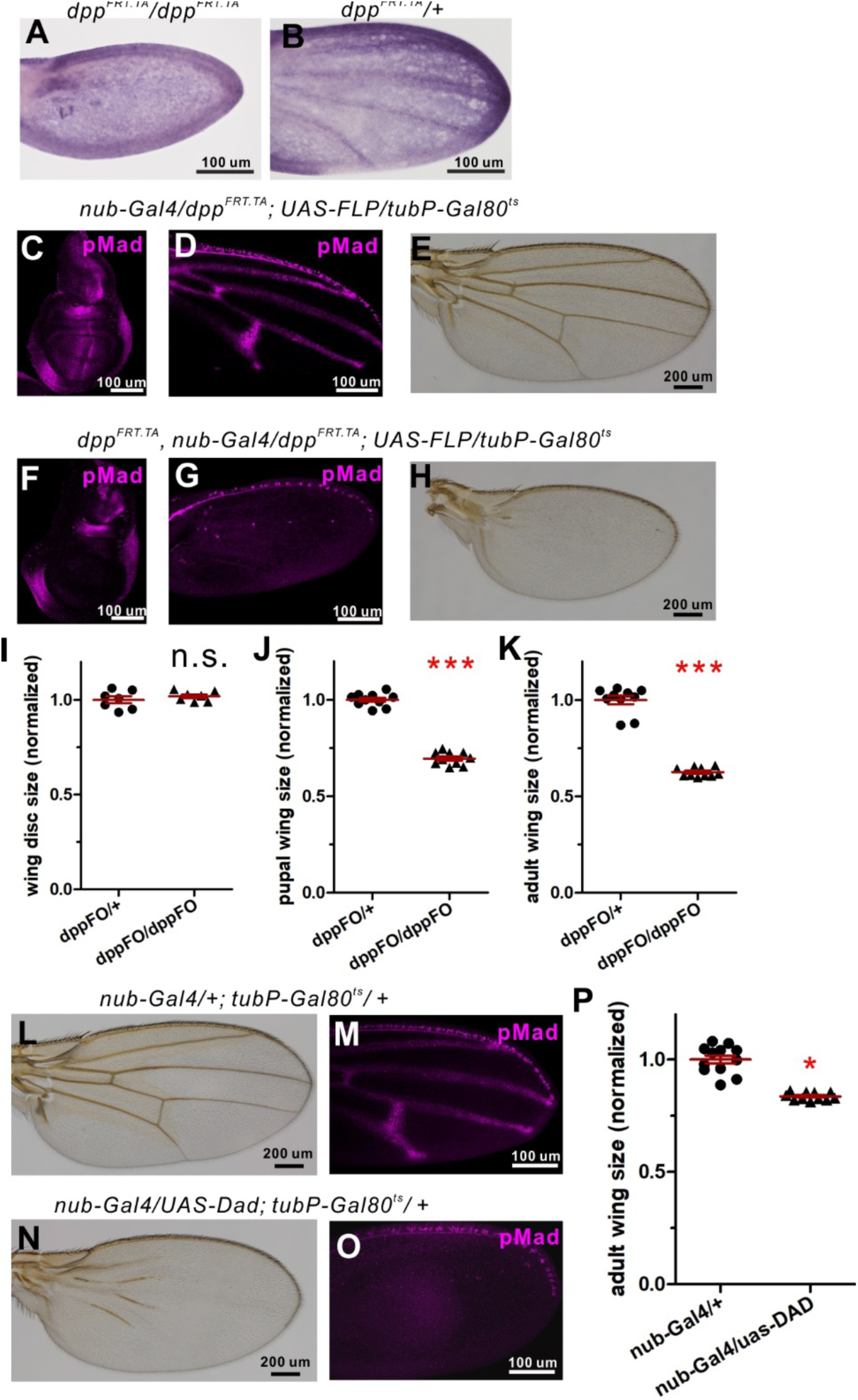
(related to Figure 1). Dpp/BMP signal regulates proliferation and patterning of the *Drosophila* pupal wing. (A, B) in situ hybridization of *dpp* probe in conditional knockout (*dpp*^*FRT.TA*/^*dpp*^*FRT.TA*^) (A) and control (*dpp*^*FRT.TA*/+)^ (B) 24h AP pupal wing. (C-E) pMad staining pattern in control (*dpp*^*FRT.TA*^*/+*) wing disc (C) and 24h AP pupal wing (D). An adult wing is shown in (E). (F-H) p^Mad^ staining pattern in *dpp*^*FO*^ (*dpp*^*FRT.TA*^*/dpp*^*FRT.TA*^) wing disc (F) and 24h AP pupal wing (G). An adult wing is shown in (H). (I-K) Size comparison between control and *dpp*^*FO*^ wingsof wing imaginal discs (I), 24h AP pupal wings (J) and adult wings (K). Means ± SEM, n.s. (not significant), ***P < 0.001, two-paired *t*-test with 95 % confidence intervals (CIs). Larvae were reared at 18°C and then transferred to 29°C 24h before pupariation, followed by dissecting wing imaginal discs (C, F and I), collecting at 24h AP and dissecting pupal (D, G, and J) or adult stage wings (E, H, and K). Sample sizes are 7 (control) and 7 (*dpp*^*FO*^) in (I), 10 (control) and 10 (*dpp*^*FO*^) in (J), and 10 (control) and 10 (*dpp*^*FO*^) in (K). (L, N) Adult wings in control (L), and *Dad* overexpression (N). (M, O) pMad expression of 24h AP pupal wings in control (M), and *Dad* overexpression (O). (P) Size comparison between control and *Dad* overexpressing wings in (L, N). Mean ± SEM. *P < 0.05, two-paired *t*-test with 95 % CIs. Sample sizes are 12 (control) and 11 (*Dad* overexpression) in (P). In (L) through (O), larvae were reared at 18° C and then transferred to 29°C as white prepupae.

**Figure S2.**
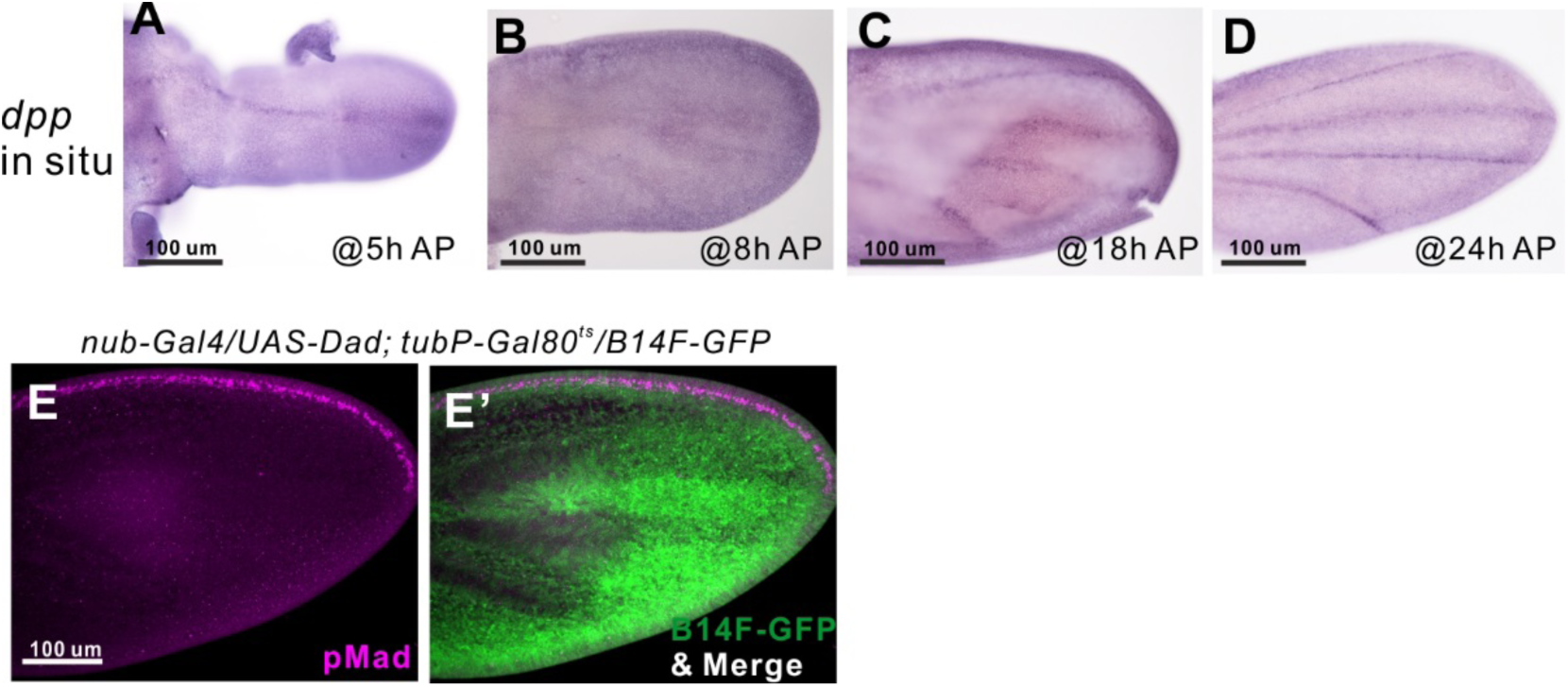
(related to Figure 2). Dpp/BMP signal forms an activity gradient in the wing tissue at 18 h AP. (A-D) in situ hybridization of *dpp* of pupal wings at 5h (A), 8h (B), 18h (C) and 24h AP (D). (E, E’) pMad expression (E) is lost and *brk*^*B14F*^*-GFP* (E’) is ubiquitously expressed in the pupal wing at 18h AP when inhibitory Smad *Dad* is overexpressed.

**Figure S3.**
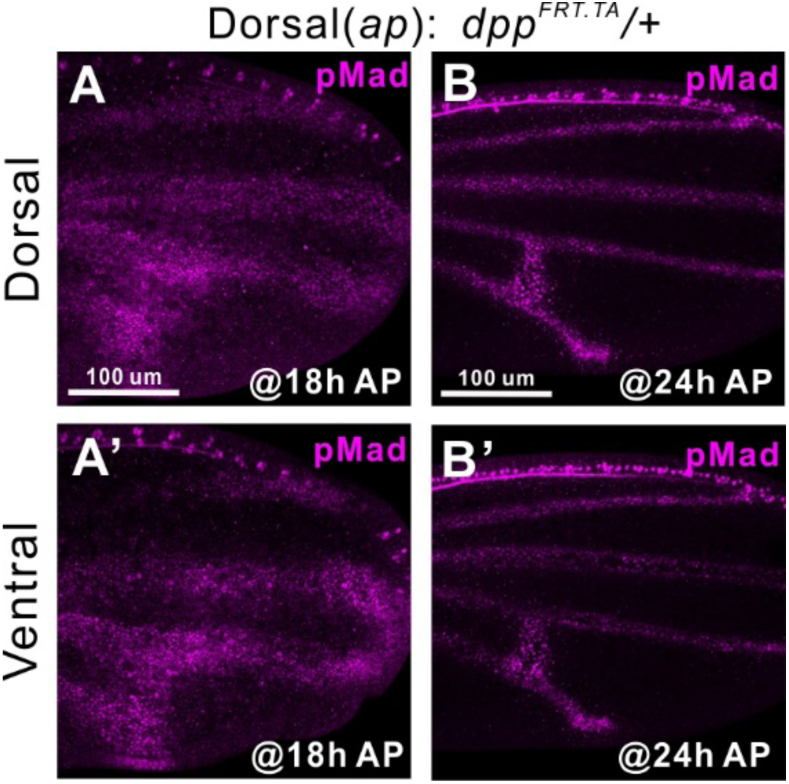
(related to Figure 3). BMP signaling between dorsal and ventral epithelia of the pupal wing. (A, B) pMad expression of dorsal (A, B) and ventral tissues (A’, B’) in control *ap>dpp*^*FO*^*/+* at 18h (A), and at 24h AP (B).

**Figure S4.**
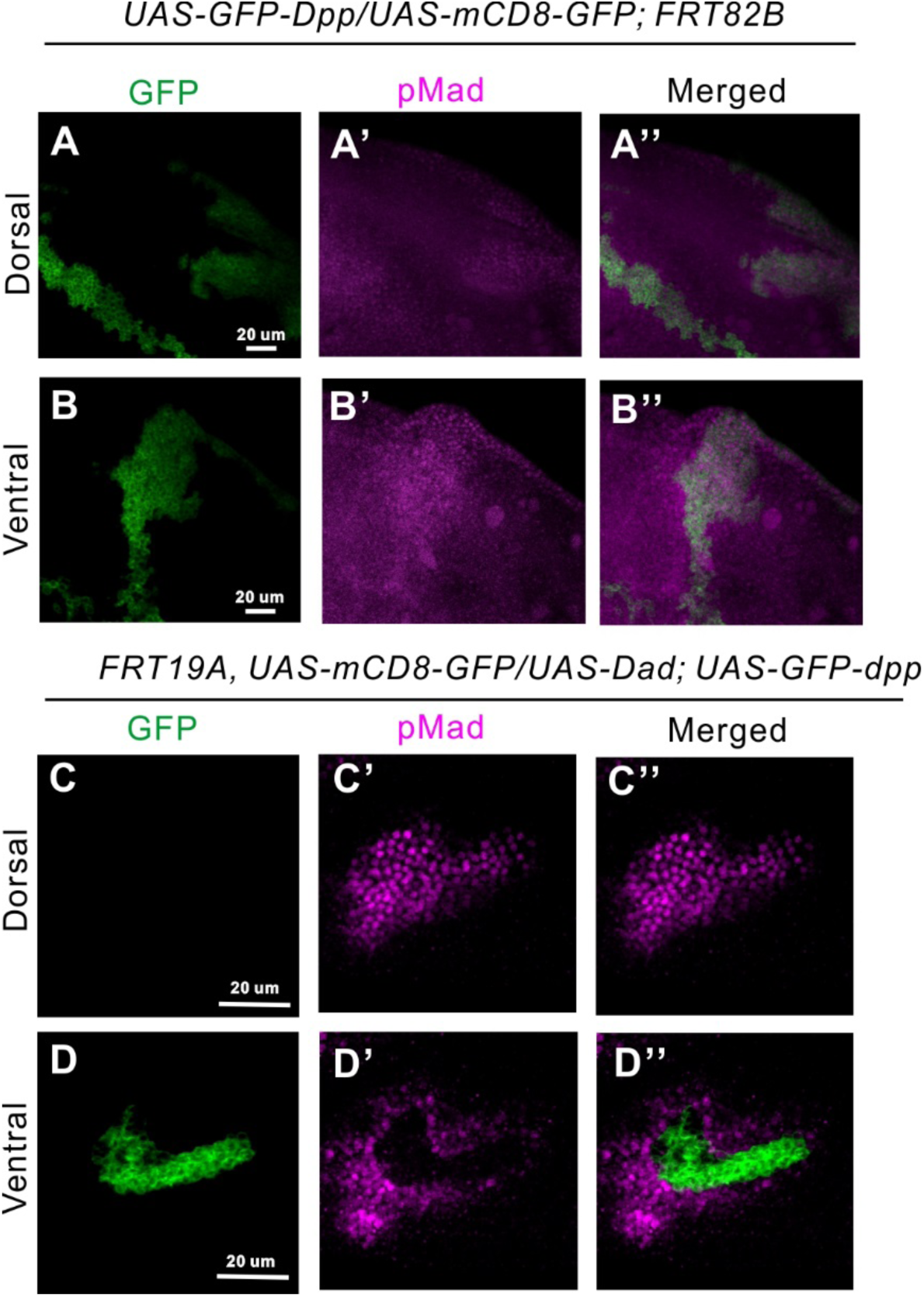
(related to Figure 4). Interplanar BMP signaling between dorsal and ventral epithelia. (A, B) Dorsal (A) and ventral (B) epithelia of the pupal wings expressing GFP-Dpp clones in dorsal layer (green, A) or ventral layer (green, B) at 18h AP. pMad expression (magenta, A’, B’). Merged images (A”, B”). (C, D) pMad expression (magenta) in dorsal (C’) and ventral (D’) epithelia of pupal wings expressing GFP-Dpp and Dad clones (green, C, D) at 24h AP. Merged images (C”, D”). Cell-non-autonomous 2D and 3D Dpp signaling in Dad-expressing clones.

**Figure S5.**
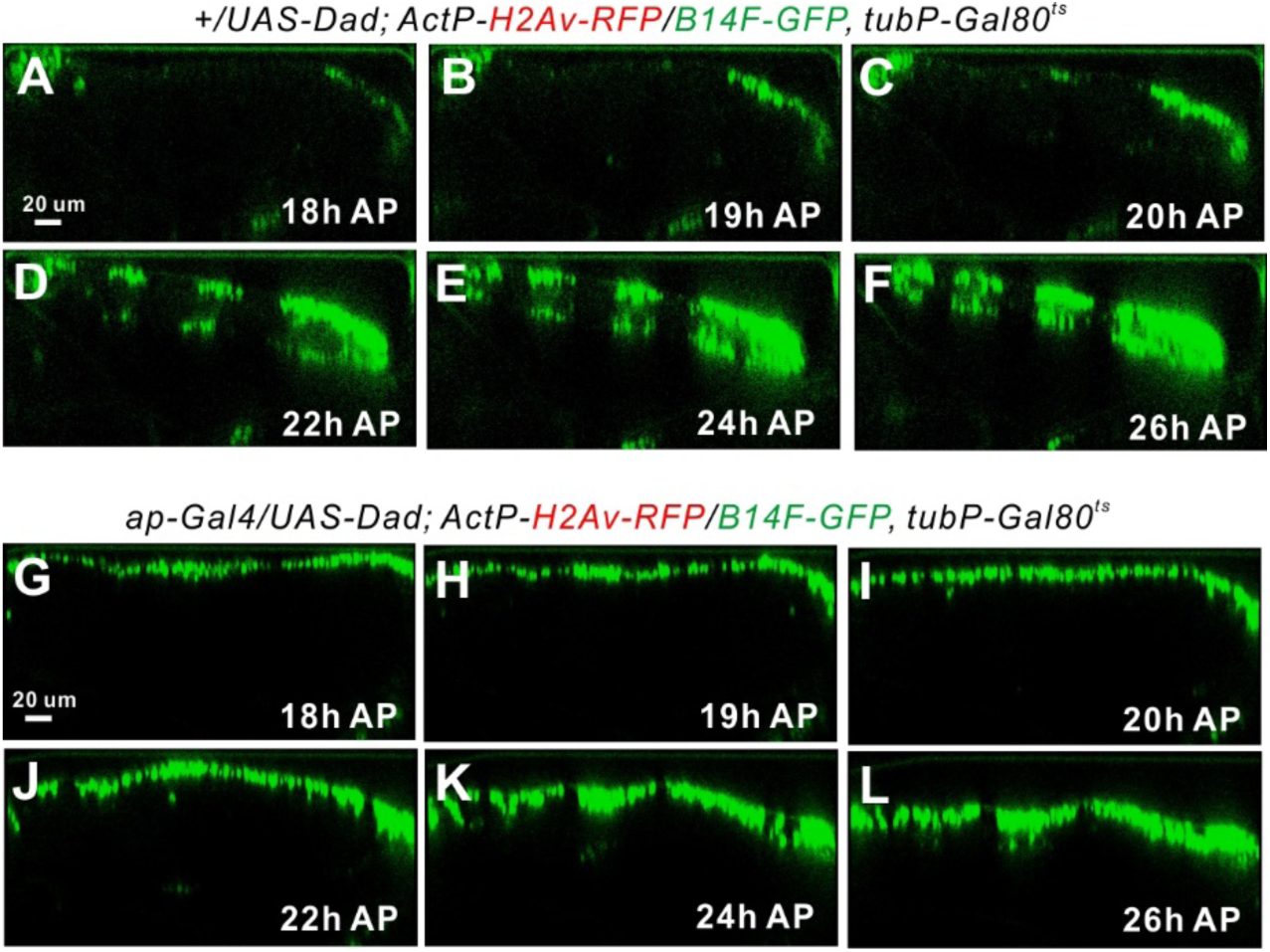
(related to Figure 5). Coupling between BMP signaling and three-dimensional tissue architecture. (A-F) Time-lapse images of anteroposterior optical cross sections of *brk*^*B14F*^*- GFP* of pupal wing at 18h (A), 19h (B), 20h (C), 22h (D) 24h (E) and 26h AP (F). (G-L) Time-lapse images of anteroposterior optical cross sections of *brk*^*B14*^*F-GFP* in Dad overexpression in the dorsal tissues of pupal wing at 18h (G), 19h (H), 20h (I), 22h (J) 24h (K) and 26h AP (L). Anterior is left and posterior right. Dorsal is up and ventral down.

**Figure S6.**
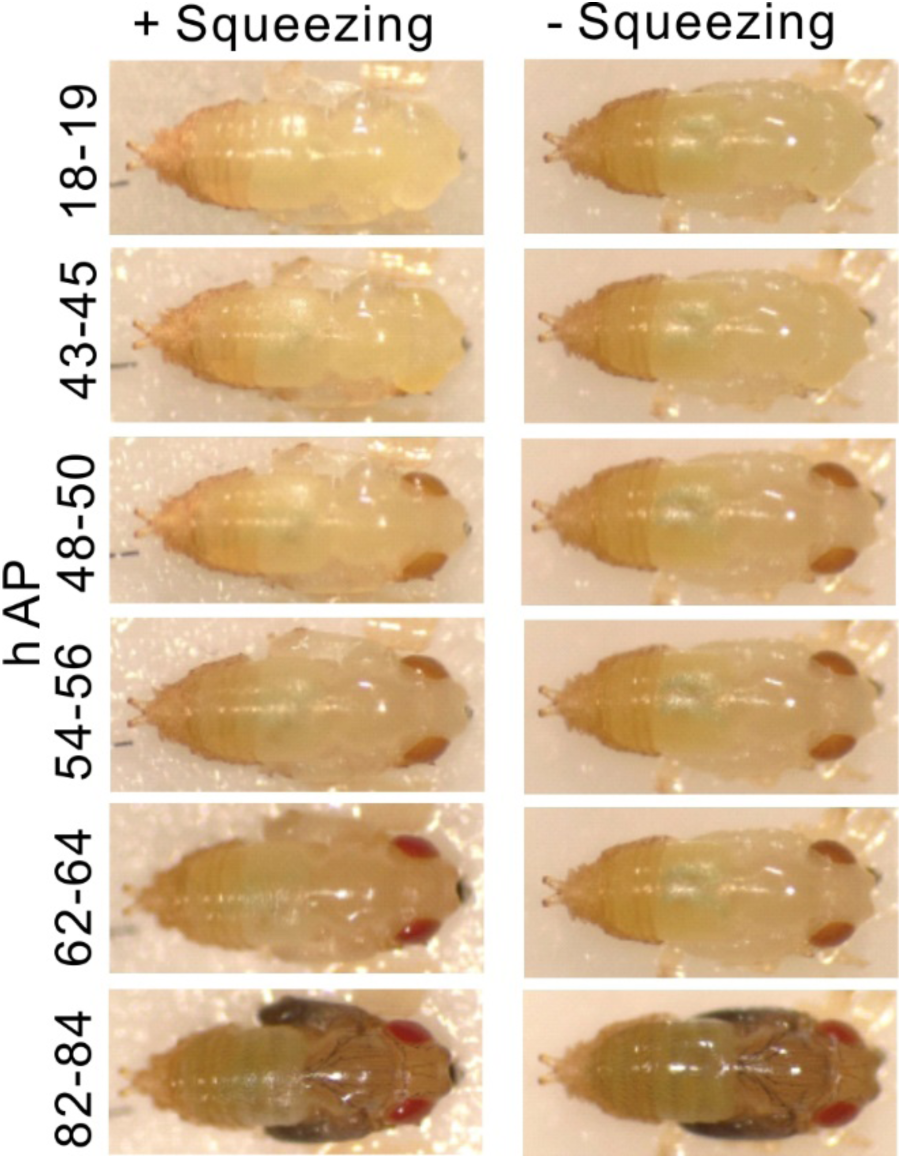
(related to Figure 6). Inducing wing inflation by abdominally squeezing pupae does not result in overall developmental delay. Once sufficient pupal case was removed to expose the pupal wings completely, the posterior, still covered in pupal case, was squeezed with forceps until the pupal wings visibly inflated. The posterior was then released and the squeezed pupa with its piece of tape was placed in a covered Petri dish with a moist Kimwipe. All pupae were then incubated at 25°C. Pupal developmental hallmarks, such as eye pigment formation, wing/cuticle darkening and scutellar bristle formation, were monitored as shown throughout the next 3-4 days. In both squeezed and unsqueezed samples, these developmental hallmarks occurred essentially in synchrony. All squeezed and unsqueezed pupae developed into live adults.

**Figure S7.**
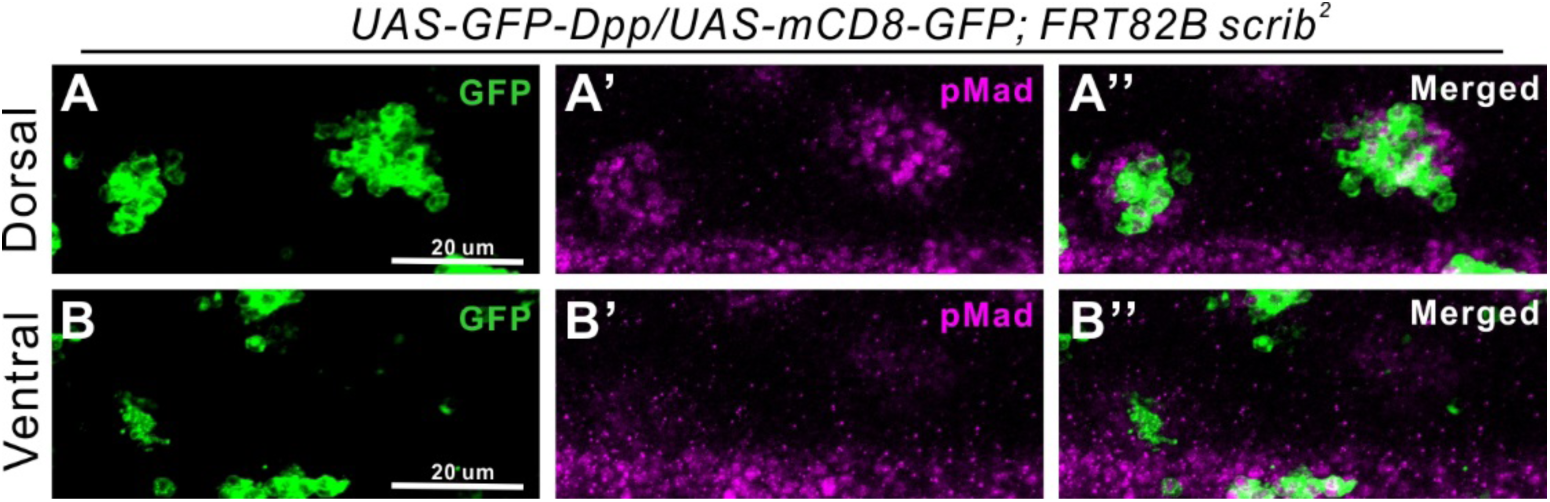
Interplanar BMP signaling between dorsal and ventral epithelia of the pupal wing. (A, B) dorsal (A) and ventral (B) epithelia of the pupal wings expressing GFP-Dpp in *scrib* clones at 24h AP (green, A, B). pMad expression (magenta, A’, B’). Merged images (A”, B”). Note that pMad staining in longitudinal vein L4 is seen below (A’, A”, B’, B”).

